# Transcriptome analysis of resistant and susceptible pepper (*Capsicum annuum* L.) in response to *Xanthomonas campestris* pv. *Vesicatoria*

**DOI:** 10.1101/2020.09.24.311274

**Authors:** Shenghua Gao, Fei Wang, Juntawong Niran, Ning Li, Yanxu Yin, Chuying Yu, Chunhai Jiao, Minghua Yao

## Abstract

Bacterial spot (BS) disease of pepper, incited by *Xanthomonas campestris* pv. *Vesicatoria* (*Xcv*), is one of the most serious diseases. For a comparative analysis of defense response to *Xcv* infection, we performed a transcriptome analysis of BS -susceptible cultivar ECW and -resistant cultivar VI037601 using the HiSeq™ 2500 sequencing platform. Approximately 140.15 G clean data were generated from eighteen libraries. From the libraries generated, we identified 52,041 genes including 35,336 reference genes, 16,705 novel transcripts, and 4,794 differentially expressed genes (DEGs). There were 1,291, 2,956, 1,795 and 2,448 DEGs in ECW-24h-vs-ECW-0h, ECW-48h-vs-ECW-0h, VI037601-24h-vs-VI037601-0h and VI037601-48h-vs-VI037601-0h groups, respectively. Interestingly, DEGs involved in disease response in the resistant variety were induced at an earlier stage and at higher levels compared with the susceptible variety. Key enriched categories included amino sugar and nucleotide sugar metabolism, sesquiterpenoid and triterpenoid biosynthesis and MAPK signaling pathway. Moreover, 273 DEGs only differentially expressed in VI037601 and 436 overlapping DEGs in ECW and VI037601 post *Xcv* inoculation, including NBS-LRR genes, oxidoreductase gene, WRKY and NAC transcription factors were identified, which were mainly involved in metabolic process, response to stimulus and biological regulation pathways. Quantitative RT-PCR of sixteen selected DEGs further validated the RNA-seq differential gene expression analysis. Our results will provide a valuable resource for understanding the molecular mechanisms of pepper resistance to *Xcv* infection and improving pepper resistance cultivars against *Xcv*.

## Introduction

Bell pepper (*Capsicum annuum* L.), an important member of the Solanaceae family, is one of the most important vegetable crops in China and many other countries [1]. It is rich in antioxidant compounds, which are essential for human health, such as capsanthin and capsaicin (Mazourek et al., 2009). In the past few decades, many research efforts have been carried out to increase pepper production because of its high nutritional and commercial value. However, pepper production has not achieved its potential yield due to biotic stresses like bacterial spot disease and anthracnose, and abiotic stresses like drought, salinity and heat [2-4]. Thus, taking more rigorous steps towards pepper productivity improvement is much needed.

Bacterial spot disease (BS), caused by gram-negative plant pathogenic bacterium *Xanthomonas campestris* pv. *vesicatoria* (*Xcv*), is a severe disease of pepper, resulting in yield and quality reduction in many pepper production areas, especially during periods of high temperatures and high moisture [5,6]. BS infection usually leads to dark lesions on the foliage and fruit of plants. Besides, lesions coalescence and leaf death could occur in severe cases [7]. The occurrence of BS has been reported all over the world, such as the USA, Northwestern Nigeria and Saudi Arabia [8-11]. BS has also occurred in China and has become more and more serious in recent years, especially in southern China. The method for controlling BS relies upon an integrated approach, which includes intensive copper-based bactericides application, crop rotation strategies, seed treatment and use of resistant cultivars. However, the most cost-effective and environmentally sustainable solution is to use resistant varieties. Because the copper-tolerance of *Xanthomonas* strains is continuously enhanced, it is particularly necessary to breed disease-resistant varieties [11].

To develop BS resistant commercially valuable cultivars through molecular breeding, four non-allelic hypersensitivity resistance genes, *Bs1*-*Bs4*, and two recessive non-hypersensitive resistance genes, *Bs5* and *Bs6*, have been identified in pepper over the years [8,12]. *Bs2, Bs3* and *Bs4* are cloned, *Bs2* and *Bs3* develop molecular markers for marker-assisted selection (MAS) [5,13-16]. *Bs5* is positioned and its linked marker is also obtained [12]. Each of these single genes described above individually confers resistance to several races of *Xcv*. For example, the *Bs2* gene confers resistance to race 0, 1, 2, 7 and 8 of *Xcv* [7]. However, each resistance gene can be overcome by specific races of the pathogen(s) in field-grown plants. Therefore, a deeper understanding of the responses of plant hosts to bacterial infection in pepper will contribute to accelerating the molecular breeding process and tackling the issue of the possible evolution of BS pathogen (s).

Plant disease resistance mechanisms are complex and rigorous for sensing and adapting to the environmental changes through the genetic regulatory network. Gene expression regulation at the transcriptional level is one important part of genetic regulations, which results in changes in signal transduction, protein synthesis and other metabolic processes [17]. Transcriptomic studies contribute to understanding the responses to bacteria. Microarrays, EST sequencing and RNA sequencing (RNA-seq) have been applied to transcriptomics analysis. RNA-seq is a powerful technology for examining the quantity and sequences of RNA using next-generation sequencing (NGS), which has been widely used to study global expression profiles and reveal DEGs involved in resistance networks under stress [18]. Besides, several transcriptome studies have been undertaken to identify relative genes involving in multiple biological processes such as fruit development, biotic and abiotic stress resistance [19-22]. In the case of BS stress mechanism, several transcriptomic studies have also been performed using microarrays and RNA-seq technique in tomato [23,24]. However, a genome-wide and comprehensive analysis of genes involved in BS resistance is not yet available in pepper.

In this study, to get deeper insights into the BS resistance mechanisms in pepper as well as to understand molecular interactions between *Xcv* and pepper, a comprehensive transcriptomic study was conducted in two pepper cultivars (ECW and VI037601) at 24h and 48h post *Xcv* strains 23-1 inoculation. Besides, we identified differentially expressed genes (DEGs), including transcription factors (TFs), kinase responsive genes, oxidoreductase and clusters of genes involved in different gene ontology terms (GO) and Kyoto encyclopedia of genes and genomes (KEGG) pathways through a comprehensive and integrated analysis of these different datasets. Furthermore, 16 induced genes were selected to further test their expression level by qRT-PCR. The transcriptome analysis will provide a subset of potential candidate defense-related genes and a valuable resource for molecular studies of pepper resistance against *Xcv* infection, which sets a good foundation for pepper disease resistance research.

## Materials and Methods

### Plant materials and pathogen inoculation

Two bell pepper genotypes, VI037601 (containing the R gene *Bs1*) and Early Calwonder (ECW) for bacterial spot resistance and susceptibility, were used for transcriptomic analysis provided by World Vegetable Center, Thailand (https://avrdc.org/). Plants were grown under standard glasshouse conditions for 16 h lighting at 25 °C/ 8 h darkness at 20 °C in a relative humidity of approximately 60%. *Xanthomonas campestris* pv. *Vesicatoria* (*Xcv*) strain 23-1 was grown at 28 °C. on nutrient agar medium as described previously [25]. Then, 2×10^8^ cfu/ml suspension of the appropriate bacterial strain *Xcv* 23-1 was mock-inoculated on the abaxial leaf surface near the midrib of the third to fifth pepper leaves, when plants were at the five leaves stage with the method as Balaji et al., described previously [23]. A water-soaked area of leaf tissue was about 1.5 to 2 cm in diameter. 2 cm leaf fragments near the bacterial infection sites, pepper leaf tissues next to the infiltration sites were collected for RNA isolation at 24h, and 48h post *Xcv* inoculation. Three leaves of the pepper with sterile water inoculation from the individual plants (ECW-0 h and VI037601-0 h) were collected as controls. Five plants were used for each sample and all fifteen collection leaves were mixed immediately, flash-frozen in liquid nitrogen and stored at -80 °C. until further use. We designated three replicates of ECW and VI037601 at 0, 24 and 48 hpi as ECW-0h-(1, 2, and 3), ECW-24h-(1, 2, and 3), ECW-48h-(1, 2, and 3), VI037601-0h-(1, 2, and 3), VI037601-24h-(1, 2, and 3) and VI037601-48h-(1, 2, and 3), respectively.

### RNA extraction, library construction and transcriptome sequencing

Total RNA was extracted from leaves of bell pepper VI037601 and ECW at different time points after inoculation using the Trizol Reagent (Life Technologies, California, USA) according to the manufacturer’s instructions, and then treated with TURBO DNase I (Promega, Beijing, China) to remove genomic DNA contamination. The integrity and concentration of all 18 RNA samples were examined by the 2100 Bioanalyzer (Agilent Technologies, Inc., Santa Clara, CA, USA) and 1.2 % agarose gel electrophoresis. The 18 prepared total RNA samples were sent to Frasergen Bioinformatics Co., Ltd (Wuhan, China) where the cDNA library was constructed using NEBNext® Ultra™ RNA Library Prep Kit for Illumina® (NEB, E7530) according to the manufacturer’s instructions. In brief, the first-strand and the second-strand cDNA were synthesized using approximately 250∼300 bp RNA inserts, which were fragmented by the enriched mRNA. After end-repair/dA-tail and adaptor ligation, the suitable fragments of double-strand cDNA were isolated by Agencourt AMPure XP beads (Beckman Coulter, Inc.), and then enriched by PCR amplification. Finally, the purity and quality of the libraries were measured by Agilent 2100 Bioanalyzer and Qubit 2.0. The eighteen cDNA libraries prepared were sequenced by Biomarker Technologies (Wuhan, China) using the Illumina HiSeq 2500 platform with pair-end (SE) 150 nt. RNA-seq was performed as previously described [21]. The raw sequence data were deposited in the Short Read Archive (SRA) National Center for Biotechnology (NCBI) Information Sequence Read Archive under the accession number PRJNA561463.

### Transcriptome analysis using reference genome-based reads mapping

The quality check was performed to eliminate low quality reads with the only adaptor, unknown nucleotides> 5%, or Q20< 20%. The high-quality clean reads that were filtered from the raw reads were mapped to the reference genome Capsicum.annuum. L_Zunla-1_Release_2.0 (http://peppersequence.genomics.cn) with TopHat 2.0 software [26]. Potential duplicate molecules were removed by examining aligned records from the aligners in BAM/SAM format. Fragments per kilobase of transcript per million fragments mapped read (FPKM) values were used to calculate the gene expression levels based on Cufflinks software [27].

### Functional analysis of differentially expressed genes

R package DEGseq was used to identify the differentially expressed genes (DEGs) between different samples after inoculation treatment of two phenotypes [28]. The ratio of the FPKM values was used to calculate the gene abundance differences between those samples. The false discovery rate (FDR) values were applied as standards to characterize the significance of the gene expression level. The genes with an absolute value of |log2(fold change)|≥1 and FDR significance score<0.05 were accepted to represent DEGs, which were used for further analysis.

To characterize the putative functions of DEGs, functional annotations of the DEGs were searched against the National Center for Biotechnology Information (NCBI) non-redundant protein (Nr) database, Swiss-Prot database with a cut-off E-value of 10^−5^ by BLASTX and the NCBI non-redundant nucleotide sequence (Nt) database using BLASTn by a cut-off E-value of 10^−5^. Besides, classification and enrichment by GO term of DEGs were performed with Blast2GO (version 3.0) (http://www.blast2go.com/) [29]. KEGG enrichment pathways analysis of DEGs was carried out using Cytoscape software (version 3.2.0) (http://www.cytoscape.org/) with the ClueGO plugin by a hypergeometric test and the Benjamini-Hochberg FDR correction (FDR ≤ 0.05).

### Identification of transcription factors (TFs)

Transcription factors were identified using PlantTFDB (http://planttfdb.cbi.pku.edu.cn/index.php), which included the sequences of 58 plant transcription factor families from 165 plant species [30]. The unigene sequence was compared with the transcription factor database by Blastx alignment, and the gene with the best E value less than e^-5^ was selected as the annotation information of the unigenes.

### Quantitative RT-PCR (qRT-PCR) analysis

The relative expression levels of selected different genes from DEGs were examined by qRT-PCR to validate the RNA-Seq data. The corresponding mRNA sequences of the selected genes were searched from the Sol Genomics Network (SGN) (https://www.solgenomics.net/). All primers for qRT-PCR were designed according to the transcript sequences using Primer Premier 5.0 and the primers used in this experiment were listed in Table S10. Besides, approximately 2 μg of total RNA was isolated from infected leaves of ECW and VI037601 by TRIzol reagent, which was used to synthesize the cDNA through the cDNA synthesis kit (TransGen, Beijing, China) according to the manufacturer’s instructions.

Quantitative RT-PCR (qRT-PCR) was performed in 96-well plates on Thermo Fisher Scientific Biosystems QuantStudio 5 Real-Time PCR system (Applied Biosystem, MA, USA) using SYBR Premix Ex Taq™ Kit (Takara, Dalian, China). The protocols of qRT-PCRs were used as follows: 95 °C for 5 min, followed by 40 cycles of 95 °C for 10 s, 58 °C for 20 s, and 72 °C for 15 s, plus melting curves to verify PCR products. Ubiquitin-conjugating protein CaUbi3 (Accession Number: AY486137.1) was used as an internal reference [31]. Three independent biological replicates were analyzed. The relative expression level of the selected genes was calculated with the 2^-ΔΔCT^ method [32].

## Results

### RNA sequencing of pepper leaves after Xcv infection and assembly of transcriptome

Two pepper cultivars, ECW and VI037601 (*Capsicum annuum* L.) were infected by *Xanthomonas campestris* pv. *Vesicatoria* (*Xcv*) via injection inoculation method, for the confirmation of resistant and susceptible genotypes. As expected, a hypersensitive response symptom was observed in VI037601 containing the R gene *Bs1* at 24 h or 48 h post *Xcv* infection (hpi), whereas cultivar ECW presented no signs at 24 hpi, even at 48 hpi, indicating that it might be different in transcriptome regulation between VI037601 and ECW post *Xcv* inoculation (Fig 1).

**Fig 1.**
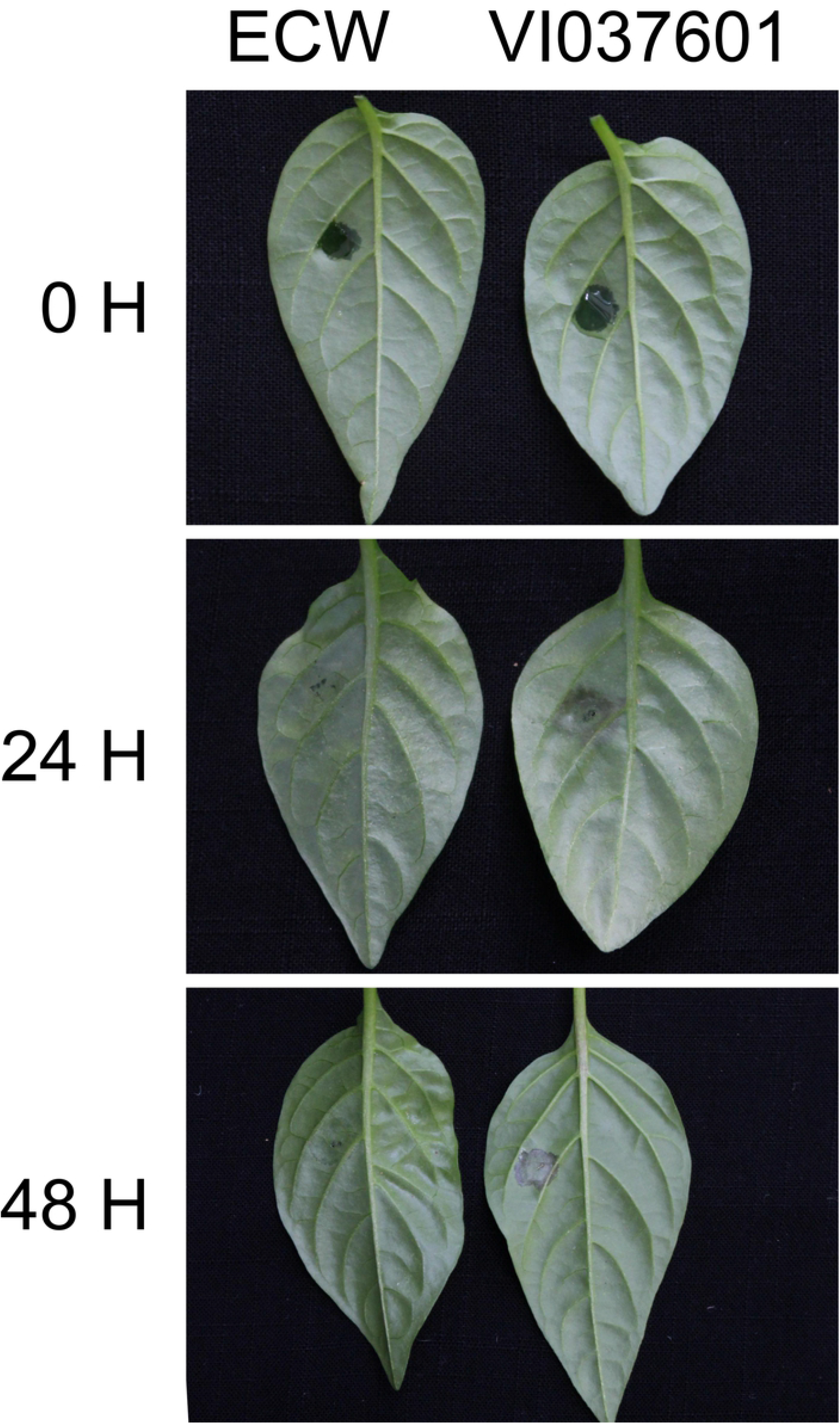
Reaction patterns of ECW and VI037601 to *Xcv* strain 23-1. 0 hpi represented mock-treatment by inoculating sterile water with a needleless syringe. 24 hpi and 48 hpi represented 24 hours and 48 hours post *Xcv* inoculation with a needleless syringe, respectively.

To accurately evaluate the comparative expression of genes in ECW and VI037601, eighteen cDNA libraries were used for RNA sequencing. Approximately 140.15 G clean data were generated using an Illumina HiSeq 2500 sequencing platform. Additional, clean data of each library was ranged from 24,362,137 to 30,342,845 clean-read pairs and GC contents of which were ranged from 42.6% to 44.2% (Table 1; Table S1). To assess the quality of the RNA-seq data, the sequence reads were mapped to the pepper reference genome Zunla-1 using TopHat2 and implementing Bowtie [33]. The number of clean reads, 71.94% -76.76% were mapped to the location within the pepper reference genome Zunla-1 (Table 1). A total of 52,043 genes, including 16,705 novel genes, were identified in this study (Table S2; Table S3). There were 38,820, 38,894, 38,719, 38,938, 39,304, and 39,080 expressed genes in ECW-0h, ECW-24h, ECW-48h, VI03760-0h, VI03760-0h, and VI03760-0h, respectively (Table S3). The high range of genome coverage of our RNA-Seq data revealed that the RNA-seq data were sufficient for subsequent bioinformatics analysis.

**Table 1.** Summary of RNA-Seq data

| Sample<sup>a</sup> | Clean reads<br>pairs | Total mapped(%) | GC content<br>(%) | Q20(%) | No. of matched<br>genes |
| --- | --- | --- | --- | --- | --- |
| ECW-0h-1 | 27,725,514 | 20,867,777(75.27) | 43.6;43.5 | 98.0;97.6 | 35,024 |
| ECW-0h-2 | 26,982,377 | 19,720,223(73.09) | 42.9;42.9 | 97.8;97.1 | 35,011 |
| ECW-0h-3 | 27,222,399 | 19,585,040(71.94) | 44.2;44.2 | 98.0;97.6 | 35,475 |
| ECW-24h-1 | 30,006,513 | 22,722,195(75.72) | 43.3;43.3 | 97.9;97.4 | 36,240 |
| ECW-24h-2 | 27,352,584 | 20,479,111(74.87) | 43.5;43.5 | 98.1;97.3 | 35,340 |
| ECW-24h-3 | 30,912,290 | 23,415,414(75.75) | 42.8;42.8 | 98.2;97.2 | 35,972 |
| ECW-48h-1 | 27,770,311 | 20,598,260(74.17) | 43.1;43.1 | 97.8;97.4 | 35,985 |
| ECW-48h-2 | 24,362,137 | 18,553,505(76.16) | 43.6;43.5 | 98.0;97.7 | 35,748 |
| ECW-48h-3 | 29,227,559 | 21,951,467(75.11) | 43.1;43.1 | 98.1;95.8 | 34,754 |
| VI037601-0h | 25,178,592 | 18,684,315(74.21) | 42.9;42.8 | 98.0;97.6 | 35,764 |
| VI037601-0h | 26,962,594 | 20,197,428(74.91) | 43.2;43.1 | 98.0;97.7 | 35,921 |
| VI037601-0h | 29,840,777 | 21,542,306(72.19) | 44.0;44.0 | 98.1;97.5 | 35,142 |
| VI037601-24 | 25,767,973 | 19,778,776(76.76) | 43.4;43.4 | 97.8;97.4 | 36,448 |
| VI037601-24 | 30,083,604 | 23,032,175(76.56) | 43.1;43.1 | 98.1;97.6 | 35,749 |
| VI037601-24 | 29,636,013 | 22,640,295(76.39) | 43.3;43.2 | 98.0;97.4 | 36,221 |
| VI037601-48 | 24,760,980 | 18,713,064(75.57) | 43.9;43.9 | 97.7;96.9 | 34,500 |
| VI037601-48 | 30,342,845 | 23,145,313(76.28) | 42.7;42.6 | 98.3;97.9 | 35,345 |
| VI037601-48 | 27,475,162 | 20,661,156(75.20) | 43.0;43.0 | 98.1;97.6 | 36,203 |

<sup>a</sup> ECW-0h and VI037601-0h: ECW and VI037601 mock-treatment with the sterile water, respectively. ECW-24h, VI037601-24h, ECW-48h and VI037601-48h: ECW and VI037601 inoculated with *Xcv*, respectively.

### Expression analysis and identification of differentially expressed genes

To investigate the expression patterns of genes in pepper leaves during the different stages after *Xcv* infection, a total of 4,794 DEGs were perceived in ECW and VI037601 at 24 hpi and 48 hpi, including 1,418 novel genes (Table S4). Among them, 1,291 (999 up regulated and 292 down regulated) and 2,956 (1,825 up regulated and 1,131 down regulated) DEGs were found at 24 hpi and 48 hpi in ECW, respectively. 1,795 (1,063 up regulated and 732 down regulated) and 2,448 (1,249 up regulated and 1,199 down regulated) DEGs were identified at 24 hpi and 48 hpi in VI037601, respectively (Fig 2A,B; Table S4). However, 1,058 (875 up regulated and 183 down regulated) and 1,054 (682 up regulated and 372 down regulated) DEGs overlapped at 24 hpi and 48 hpi in ECW and VI037601, respectively (Fig 2A; Table S4). 436 overlapping DEGs were found in ECW and VI037601 post *Xcv* inoculation (Fig 2A; Table S5). Furthermore, 273 DEGs were specific differentially expressed in VI037601 post *Xcv* inoculation (Fig 2A; Table S5). Overall, the number of DEGs was considerably higher at 48 hpi than that at 24 hpi in ECW and VI037601, the number of up-regulated DEGs was greater than that of down-regulated DEGs in ECW and VI037601 at 24 hpi and in ECW at 48 hpi. Besides, the number of DEGs at 24 hpi was higher in VI037601 than that at 24 hpi in ECW, especially down regulated DEGs. In contrast, less DEGs were identified at 48 hpi in VI037601 compared with that at 48 hpi in ECW, especially up regulated DEGs, whereas the number of down regulated DEGs were almost the same (Fig 2B). These results indicated that more DEGs in VI037601 were involved in the response to *Xcv* inoculation in the early stages compared with ECW. The DEGs might contain the disease resistance gene(s), such as *Bs1*, which conferred resistance to *Xcv* in VI037601.

**Fig 2.**
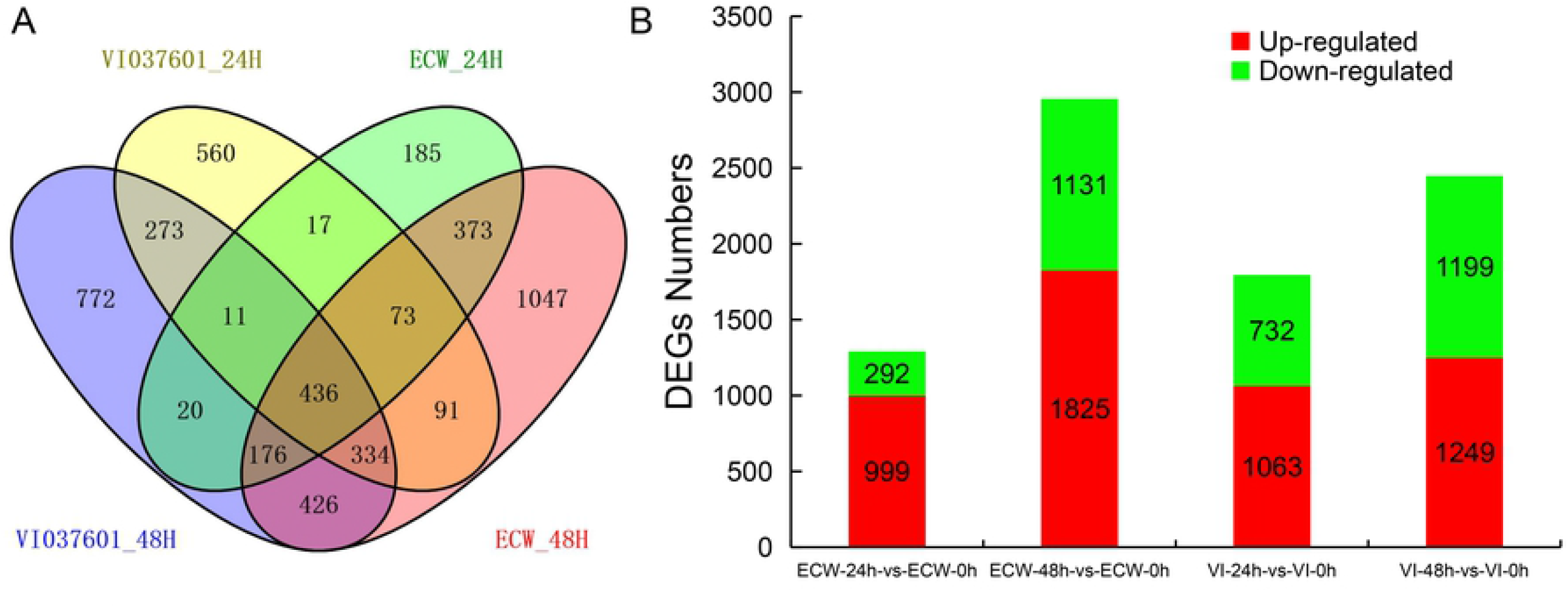
Expressional analysis of DEGs in ECW and VI037601 leaves at 24 hours and 48 hours post *Xcv* inoculation with *Xcv*. (**A**) Numbers of DEGs at 24 hpi and 48 hpi in ECW and VI037601, or between ECW and VI037601 at different time points. (**B**) Numbers of up- and down-regulated DEGs at 24 hpi and 48 hpi in ECW and VI037601, respectively.

### Functional enrichment analysis of DEGs

To acknowledge the putative functions and pathways of the DEGs in *Xcv* inoculation transcriptomes among the different stages in ECW and VI037601, GO classification and KEGG functional enrichment analyses of DEGs were performed by GOseq and KOBAS 2.0 [34-36]. In total, 4,794 DEGs in ECW-24h-vs-ECW-0h, ECW-48h-vs-ECW-0h, VI037601-24h-vs-VI037601-0h and VI037601-48h-vs-VI037601-0h groups were annotated with GO terms and assigned to three categories: biological process (BP), cellular component (CC), and molecular function (MF). The most enriched GO terms in BP categories included single-organism metabolic process (GO:0044710), oxidation-reduction process (GO:0055114), response to stress (GO:0006950), defense response (GO:0006952), and secondary metabolic process (GO:0019748) in four comparison groups (Table S6). Oxidoreductase activity (GO:0016491), peroxidase activity (GO:0004601), oxidoreductase activity, acting on peroxide as acceptor (GO:0016684), antioxidant activity (GO:0016209), chitinase activity (GO:0004568), and chitin binding (GO:0008061) were enriched in the MF category. Furthermore, extracellular region (GO:0005576) were enriched in CC category (Table S6). However, regulation of protein serine/threonine phosphatase activity (GO:0080163) in BP category, UDP-glucosyltransferase activity (GO:0035251), transferase activity, transferring hexosyl groups (GO:0016758), regulation of protein serine/threonine phosphatase activity (GO:0080163), and hormone binding (GO:0042562) in MP category were found to be differentially expressed in VI037601-24h-vs-VI037601-0h and VI037601-48h-vs-VI037601-0h (Fig. 3; Table S6). These processes associated with disease resistance were enriched, indicating that the corresponding genes of these significant terms might play important roles in resistance to *Xcv* inoculation.

**Fig 3.**
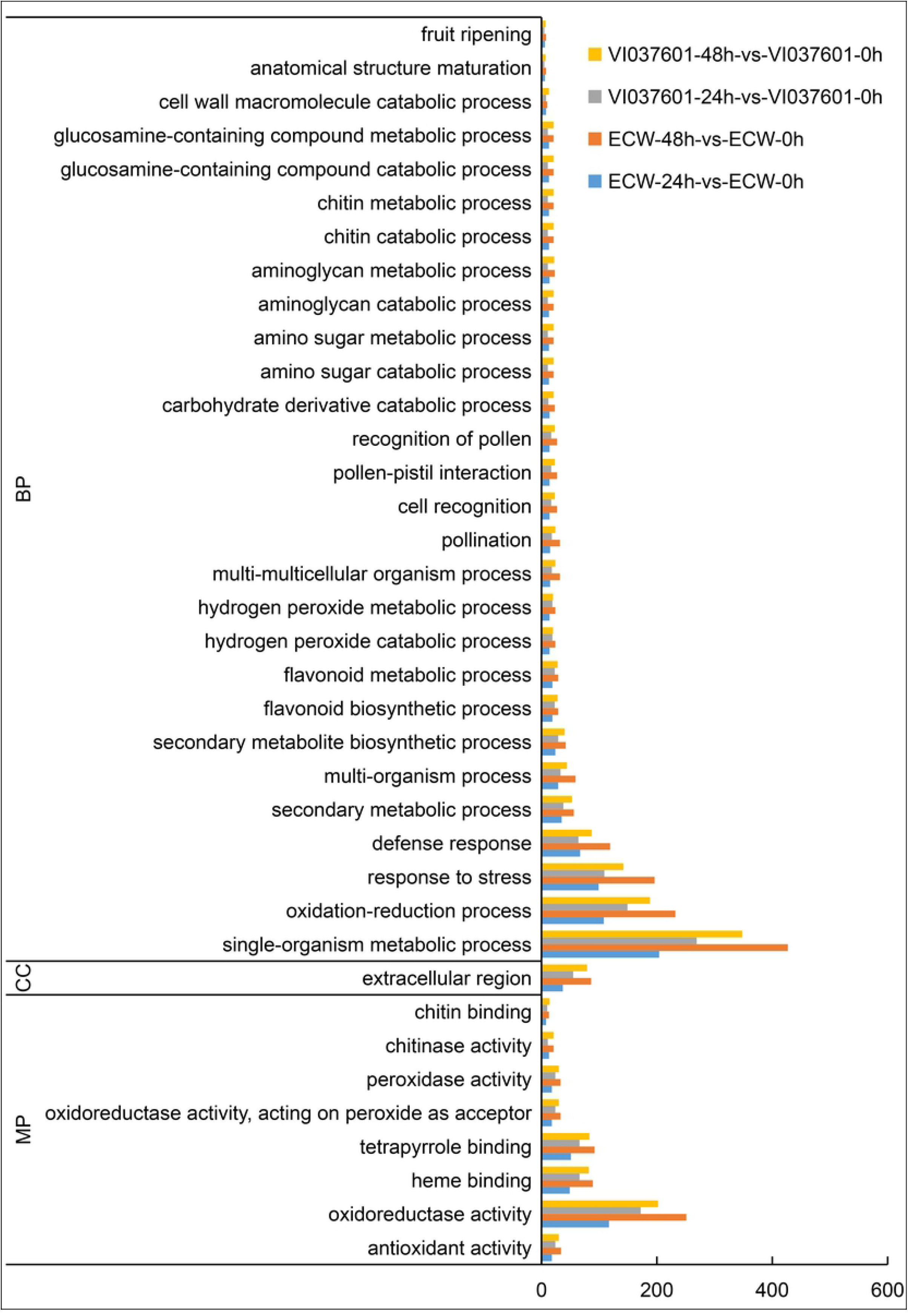
GO enrichment of DEGs in ECW and VI037601 post inoculation.

The significant KEGG enrichment pathways categories in the four comparisons were represented in this study. DEGs were largely enriched in Amino sugar and nucleotide sugar metabolism (ko00520), Sesquiterpenoid and triterpenoid biosynthesis (ko00909) and MAPK signaling pathway (ko04016) in the four comparisons (Table 2; Table S7). Fatty acid elongation (ko00062), phenylalanine metabolism (ko00360), phenylalanine, tyrosine and tryptophan biosynthesis (ko00400), glutathione metabolism (ko00480), phenylpropanoid biosynthesis (ko00940), stilbenoid, diarylheptanoid and gingerol biosynthesis (ko00945) and biosynthesis of unsaturated fatty acids (ko01040) were also enriched in the comparisons, except for the comparison ECW-24h-vs-ECW-0h (Table 2; Table S7). Besides these above-mentioned pathways, plant-pathogen interaction (ko04626) was significantly enriched with up regulated DEGs in ECW-48h-vs-ECW-0h and VI037601-48h-vs-VI037601-0h. Ubiquinone and another terpenoid-quinone biosynthesis (ko00130), flavonoid biosynthesis (ko00941), anthocyanin biosynthesis (ko00942), biosynthesis of amino acids (ko01230) and plant hormone signal transduction (ko04075) were enriched in the comparisons ECW-48h-vs-ECW-0h and VI037601-24h-vs-VI037601-0h. Yet, tyrosine metabolism (ko00350) and isoquinoline alkaloid biosynthesis (ko00950) were enriched in VI037601 after infection. These results indicated that the different expression patterns of DEGs in significant KEGG enrichment pathway categories in ECW and VI037601 helped to determine the functions of DEGs and screen of candidate resistance genes, which was responsible for the resistance to *Xcv* in VI037601.

**Table 2.** Part significantly enriched KEGG pathways of DEGs in ECW and VI037601 after *Xcv* infection.

| KEGG Pathway | Pathway ID | No. of DEGs (%) |  |  |  |
| --- | --- | --- | --- | --- | --- |
|  |  | ECW-24h | ECW-48h | VI-24h | VI-48h |
| Amino sugar and nucleotide sugar metabolism | ko00520 | 17(7.91) | 34(15.81) | 16(14.43) | 31(13.40) |
| Sesquiterpenoid and triterpenoid biosynthesis | ko00909 | 16(16.49) | 19(19.59) | 14(13.11) | 13(15.41) |
| MAPK signaling pathway- plant | ko04016 | 24(7.38) | 39(12.00) | 24(7.38) | 34(10.46) |
| Fatty acid elongation | ko00062 | - | 9(16.36) | 11(20.00) | 10(18.18) |
| Phenylalanine metabolism | ko00360 | - | 14(16.09) | 14(16.09) | 18(20.69) |
| Phenylalanine, tyrosine and tryptophan biosynthesis | ko00400 | - | 15(19.23) | 13(17.67) | 14(10.88) |
| Glutathione metabolism | ko00480 | - | 17(11.56) | 26(7.44) | 16(14.42) |
| Phenylpropanoid biosynthesis | ko00940 | - | 51(16.72) | 40(11.67) | 47(15.00) |
| Stilbenoid, diarylheptanoid and gingerol biosynthesis | ko00945 | - | 9(10.98) | 10(12.20) | 11(13.14) |
| Biosynthesis of unsaturated fatty acids | ko01040 | - | 13(21.67) | 7(13.11) | 9(10.46) |
| Plant-pathogen interaction | ko04626 | - | 51(10.6) | - | 40(8.32) |
| Ubiquinone and other terpenoid-quinone biosynthesis | ko00130 | - | 8(11.94) | 8(11.94) | - |
| Flavonoid biosynthesis | ko00941 | - | 10(8.80) | 10(9.80) | - |
| Anthocyanin biosynthesis | ko00942 | - | 3(16.67) | 4(22.22) | - |
| Biosynthesis of amino acids | ko01230 | - | 36(10.06) | 23(6.42) | - |
| Plant hormone signal transduction | ko04075 | - | 34(8.08) | 39(9.26) | - |
| Tyrosine metabolism | ko00350 | - | - | 10(11.24) | 11(12.36) |
| Isoquinoline alkaloid biosynthesis | ko00950 | - | - | 8(14.8) | 9(16.67) |

ECW-24h and ECW-48h represented the comparisons ECW-24h-vs-ECW-0h and ECW-48h-vs-ECW-0h, respectively. VI-24h and VI-48h represented the comparisons VI037601-24h-vs-VI037601-0h and VI037601-48h-vs-VI037601-0h, respectively. (%) represented the percentage of DEGs in total genes of the KEGG pathway.

### Transcriptional changes in response to Xcv infection

Many genes play a critical role in recognizing pathogen-associated molecular patterns (PAMPs) and subsequently activating plant defense mechanisms in response to pathogen attacks, such as kinases, pathogenesis-related protein, oxidoreductase and E3 ubiquitin-protein ligase [21,37]. In this study, thirteen specific differentially expressed protein kinase genes and 22 specific differentially expressed oxidoreductase genes were identified in VI037601 post *Xcv* inoculation (Table S5). We also found that Capana09g000319 (aldehyde dehydrogenase) and Capana09g000326 (glycosyltransferase) were significantly differentially expressed in VI037601 post *Xcv* inoculation, but not expressed in ECW (Table S5). Moreover, 29 differentially expressed kinase responsive genes (26 up-regulated and 3 down-regulated genes), including LRR receptor-like ser/thr protein kinase, were identified in ECW and VI037601 at 24 hpi and 48 hpi (Fig 4A; Table S8). Besides that, many genes encoding other disease response proteins were also identified, such as disease resistance, TMV resistance protein, pathogenesis-related proteins, receptor-like protein and proteinase inhibitor (Fig 4A; Tables S8). Interestingly, most of these DEGs were up-regulated at different time points after *Xcv* inoculation in ECW and VI037601, except for a disease-resistant protein Capana03g001868 and a novel receptor-like protein (XLOC_034810) (Fig 4A; Table S8). Other overlapping BS disease response genes, including twenty DEGs encoding cytochrome P450, four DEGs encoding E3 ubiquitin-protein ligase, 47 DEGs encoding oxidoreductase and 8 DEGs encoding chitinase were also differentially expressed and their expression levels increased after *Xcv* infection in ECW and VI037601 (Fig 4B; Table S8). However, the expression analysis of *Bs* resistance genes showed that *bs2* (Capana09g000438) and *bs3* (Capana02g001306) were not/hardly expressed in pepper leaves before and after *Xcv* inoculation. Furthermore, the expression of *bs4* (XLOC_032013), as well as the homologs of *bs2* and *bs3*, did not change significantly (Table S3).

**Fig 4.**
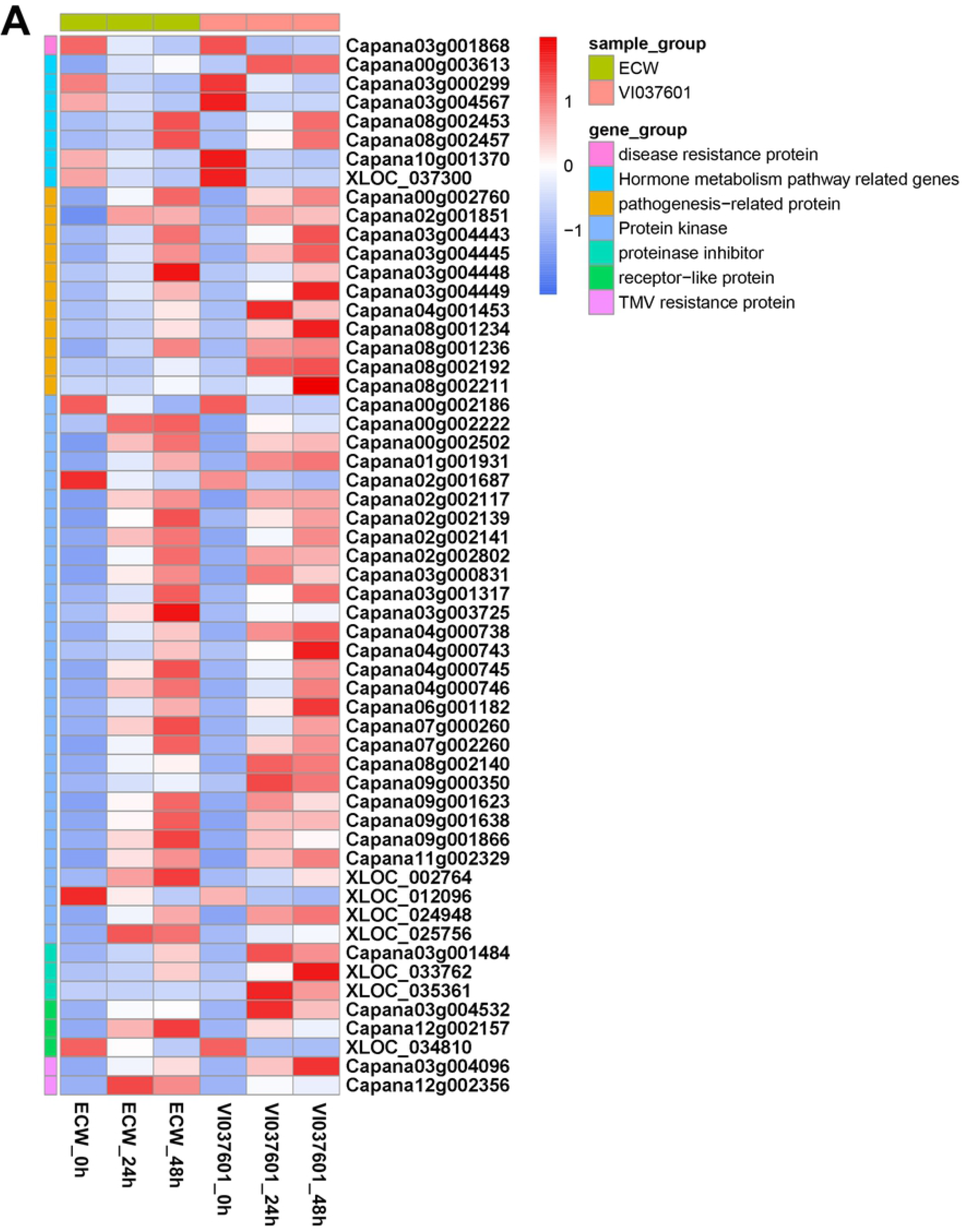
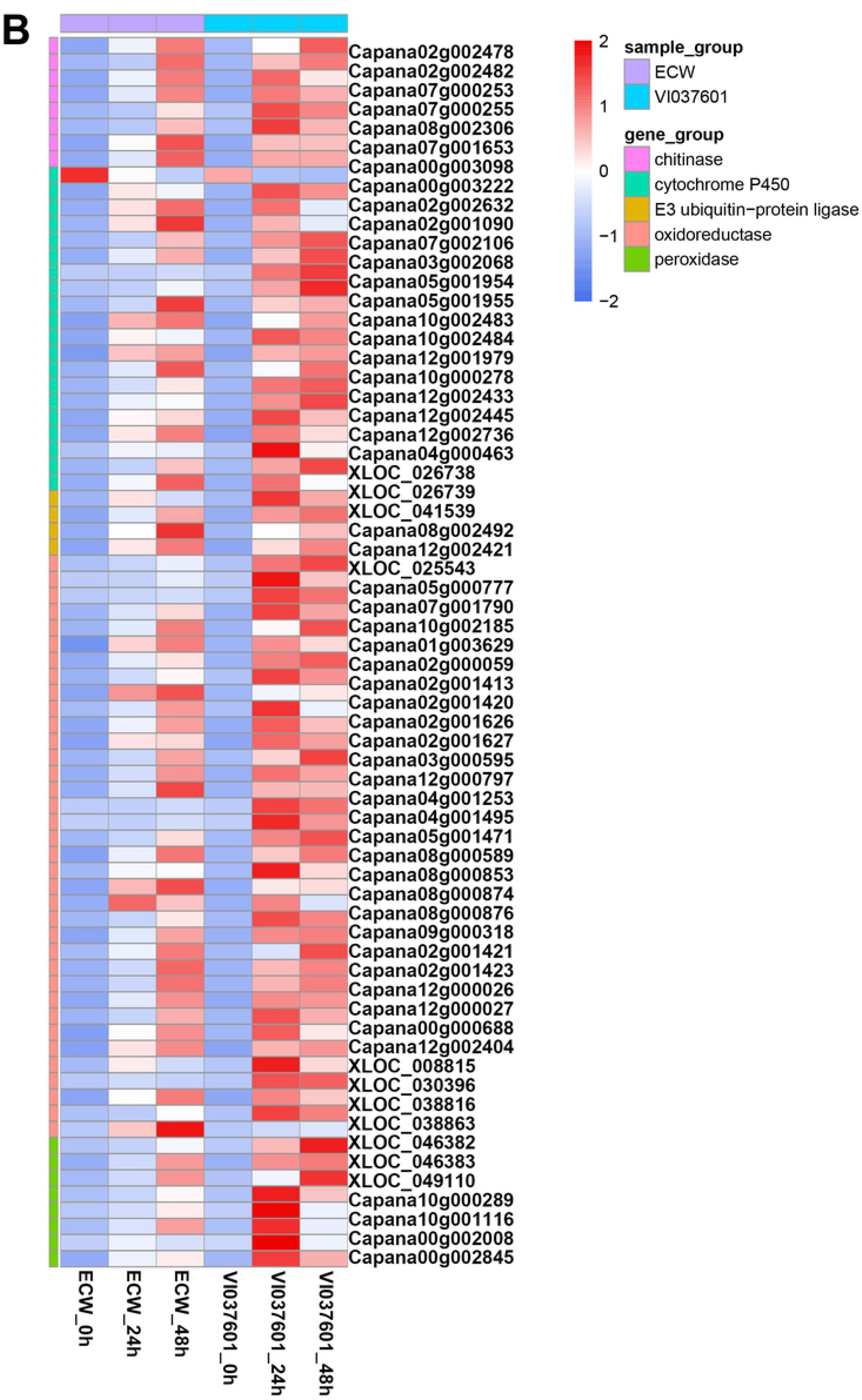
Heatmaps of the overlapping differentially expressed genes (DEGs) associated with disease resistance in ECW and VI037601 after *Xcv* inoculation. (**A**) DEGs encoding salicylic acid and auxin mediated or activated signaling pathway genes, Receptor like protein kinase, receptor-like protein, disease resistance protein, TMV resistance protein, proteinase inhibitor and pathogenesis-related protein. (**B**) DEGs encoding E3 ubiquitin-protein ligase, chitinase, peroxidase, oxidoreductase and cytochrome P450.

### The response of differentially expressed transcription factors to Xcv Infection

In plants, transcription factors (TFs) play important roles in the regulation of different physiological and biochemical programs in response to plant-pathogen interaction [21]. In our study, 254 DEGs were identified, which involved in 35 TF families, including 27 AP2-EREBP (APETALA2-ethylene-responsive element binding protein) genes, 6 heat stress transcription factor (HSF), 20 NAC, and 33 WRKY TFs (Table 3; Table S9). Among them, most DEGs were up regulated in the four groups (Table 3). Moreover, 21 TFs, including Zinc finger proteins (ZFPs), MYB, WRKY and ethylene-responsive transcription factors (ERFs) were specific differentially expressed in VI037601 post *Xcv* inoculation (Table S8). These identified TFs might be likely to perform an important role in pepper -*Xcv* interaction. Besides that, 31 putative overlapping TFs were differentially expressed in ECW and VI037601 after *Xcv* infection (Fig 5; Table S8). Among these differentially expressed TFs, all DEGs encoding MYB, WRKY, ethylene responsive factor (ERFs), HSF, and NAC TFs were up regulated after *Xcv* infection in ECW and VI037601 (Fig 5; Table S8). However, Capana05g000390 (ZFP), Capana11g001506 (bHLH79 TF) and Capana04g000966 (ZFP) were down regulated (Fig 5; Table S8).

**Table 3.** The number of DEGs identified as transcription factors in pepper.

| Category | Total | ECW_24h |  | ECW_48h |  | VI037601_24h |  | VI037601_48h |  |
| --- | --- | --- | --- | --- | --- | --- | --- | --- | --- |
|  |  | Up | Down | Up | Down | Up | Down | Up | Down |
| ABI3VP1 | 6 | 0 | 0 | 1 | 2 | 1 | 2 | 0 | 1 |
| Alfin-like | 1 | 1 | 0 | 1 | 0 | 1 | 0 | 1 | 0 |
| AP2-EREBP | 27 | 7 | 0 | 12 | 2 | 15 | 4 | 14 | 8 |
| ARF | 3 | 0 | 0 | 1 | 1 | 0 | 1 | 0 | 0 |
| BES1 | 1 | 0 | 0 | 0 | 1 | 0 | 0 | 0 | 0 |
| bHLH | 22 | 4 | 3 | 4 | 10 | 0 | 5 | 3 | 6 |
| BSD | 1 | 1 | 0 | 1 | 0 | 0 | 0 | 0 | 0 |
| bZIP | 2 | 0 | 0 | 0 | 2 | 0 | 0 | 0 | 2 |
| ZFPs | 34 | 2 | 2 | 5 | 14 | 7 | 10 | 1 | 16 |
| CSD | 1 | 0 | 0 | 0 | 0 | 0 | 0 | 0 | 0 |
| DBP | 1 | 0 | 0 | 1 | 0 | 0 | 0 | 0 | 0 |
| E2F-DP | 1 | 0 | 0 | 1 | 0 | 0 | 0 | 0 | 0 |
| FAR1 | 1 | 0 | 0 | 0 | 1 | 0 | 0 | 0 | 0 |
| FHA | 2 | 1 | 0 | 1 | 0 | 0 | 0 | 1 | 0 |
| G2-like | 5 | 0 | 0 | 2 | 1 | 2 | 2 | 1 | 0 |
| GRAS | 7 | 0 | 0 | 1 | 2 | 1 | 1 | 3 | 1 |
| GRF | 2 | 0 | 0 | 0 | 1 | 0 | 1 | 0 | 1 |
| HSF | 6 | 4 | 0 | 5 | 0 | 3 | 0 | 3 | 0 |
| LIM | 2 | 0 | 0 | 0 | 2 | 0 | 0 | 0 | 0 |
| LOB | 4 | 1 | 0 | 2 | 1 | 2 | 0 | 3 | 0 |
| MYB | 43 | 5 | 2 | 9 | 14 | 7 | 14 | 10 | 13 |
| NAC | 20 | 11 | 0 | 15 | 0 | 11 | 1 | 9 | 3 |
| OFP | 2 | 0 | 0 | 1 | 0 | 0 | 0 | 0 | 1 |
| PLATZ | 4 | 1 | 0 | 1 | 0 | 3 | 0 | 4 | 0 |
| SBP | 4 | 0 | 1 | 0 | 3 | 0 | 2 | 1 | 2 |
| SRS | 1 | 0 | 0 | 1 | 0 | 0 | 0 | 1 | 0 |
| TCP | 5 | 0 | 1 | 0 | 3 | 0 | 0 | 0 | 4 |
| Trihelix | 2 | 0 | 0 | 1 | 0 | 1 | 1 | 0 | 1 |
| TUB | 1 | 0 | 0 | 0 | 1 | 0 | 0 | 0 | 1 |
| WRKY | 33 | 22 | 0 | 28 | 0 | 15 | 1 | 17 | 1 |
| ARR-B | 1 | 0 | 0 | 0 | 0 | 0 | 0 | 0 | 1 |
| MADS | 4 | 0 | 0 | 0 | 0 | 0 | 2 | 0 | 3 |
| Tify | 2 | 0 | 0 | 0 | 0 | 2 | 0 | 1 | 0 |
| ZF-HD | 3 | 0 | 0 | 0 | 0 | 0 | 2 | 0 | 1 |

**Fig 5.**
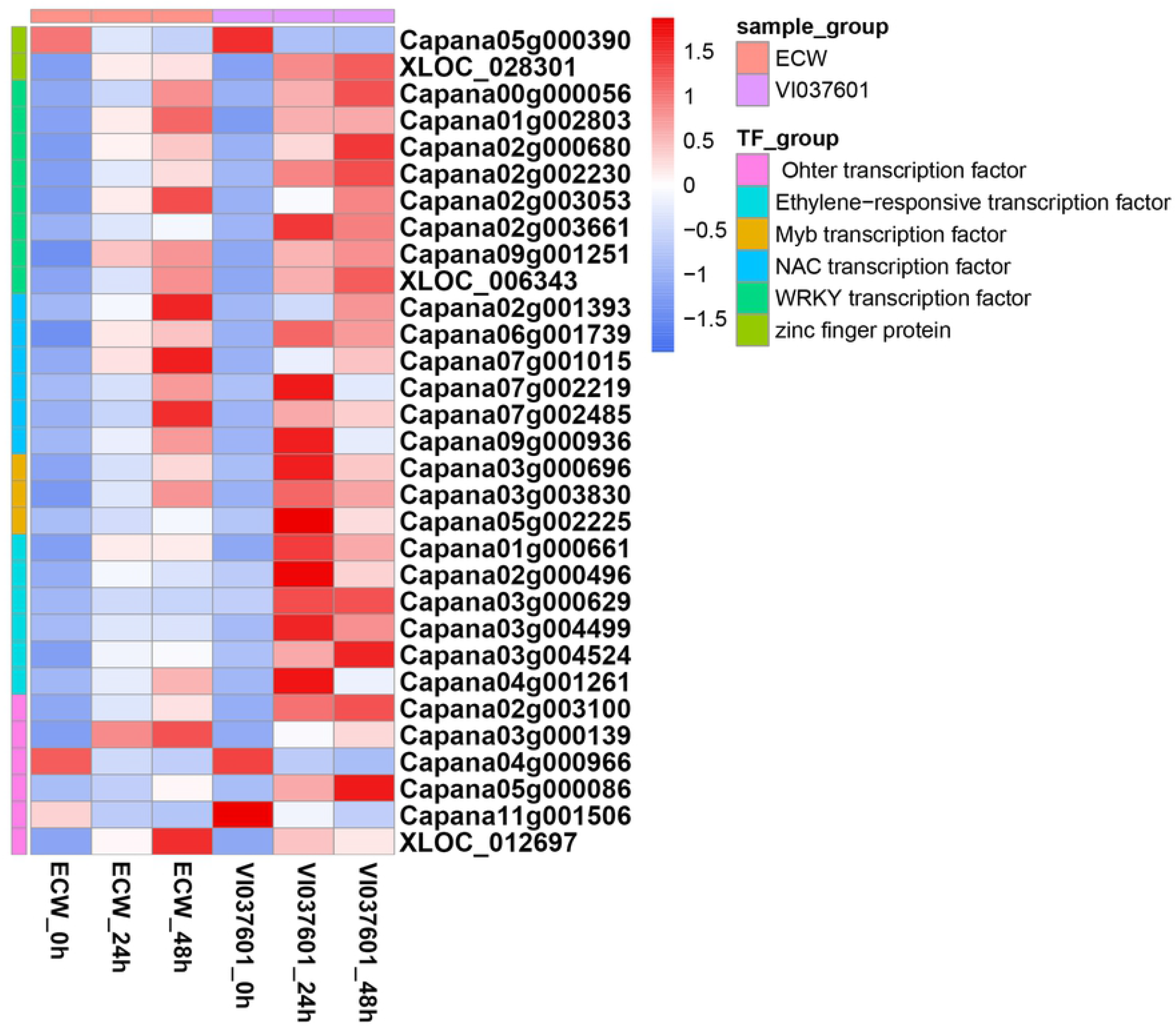
Heatmaps of the overlapping differentially expressed transcription factor genes (DEGs) in ECW and VI037601 post *Xcv* inoculation.

In brief, 28 TF responsive genes were up regulated and 3 were down regulated in ECW and VI037601 after *Xcv* infection (Fig 5; Table S8). Thus, it is most possible to defend against *Xcv* infection mainly by common up regulated genes in ECW and VI037601. Nevertheless, different down regulated DEGs might also play an important role by negatively regulating the pepper immunity upon *Xcv* infection in ECW and VI037601.

### Validation of RNA-seq data by qRT-PCR

To verify the validity of RNA-seq analysis results, transcriptional levels of 16 randomly selected DEGs representing a wide range of expression levels and patterns were detected in ECW and VI037601 post *Xcv* inoculation by qRT-PCR analysis (Fig 6). Among these 16 selected genes, majority of these DEGs were associated with massive defense response processes including receptor kinase (Capana01g001931 and Capana09g001638), protein kinase (Capana00g002502 and Capana03g000831), pathogenesis-related genes (Capana03g004445 and Capana04g001453), ERF (Capana01g000661), MYB TF (Capana05g002225), NAC TF (Capana07g001015), WRKY TF (Capana09g001251 and Capana00g000056), zinc finger protein transcription factor (Capana05g000390), disease resistance protein (Capana03g001868), and second metabolite biosynthesis (Capana01g001748, Capana04g000463 and Capana10g002483) (Fig 6). As expected, the transcription levels of the almost genes were significantly up-regulated, whereas Capana03g001868 (disease resistance protein) and Capana05g000390 (zinc finger protein) were down regulated after *Xcv* infection in ECW and VI037601. These findings showed that the expression data provided by qRT-PCR were following the profiles detected by RNA-seq at all time points in ECW and VI037601. Besides, the difference in the magnitude of DEGs identified in this study may be due to differences in algorithms of qRT-PCR and RNA-seq (Fig 4; Fig 5; Fig 6).

**Fig 6.**
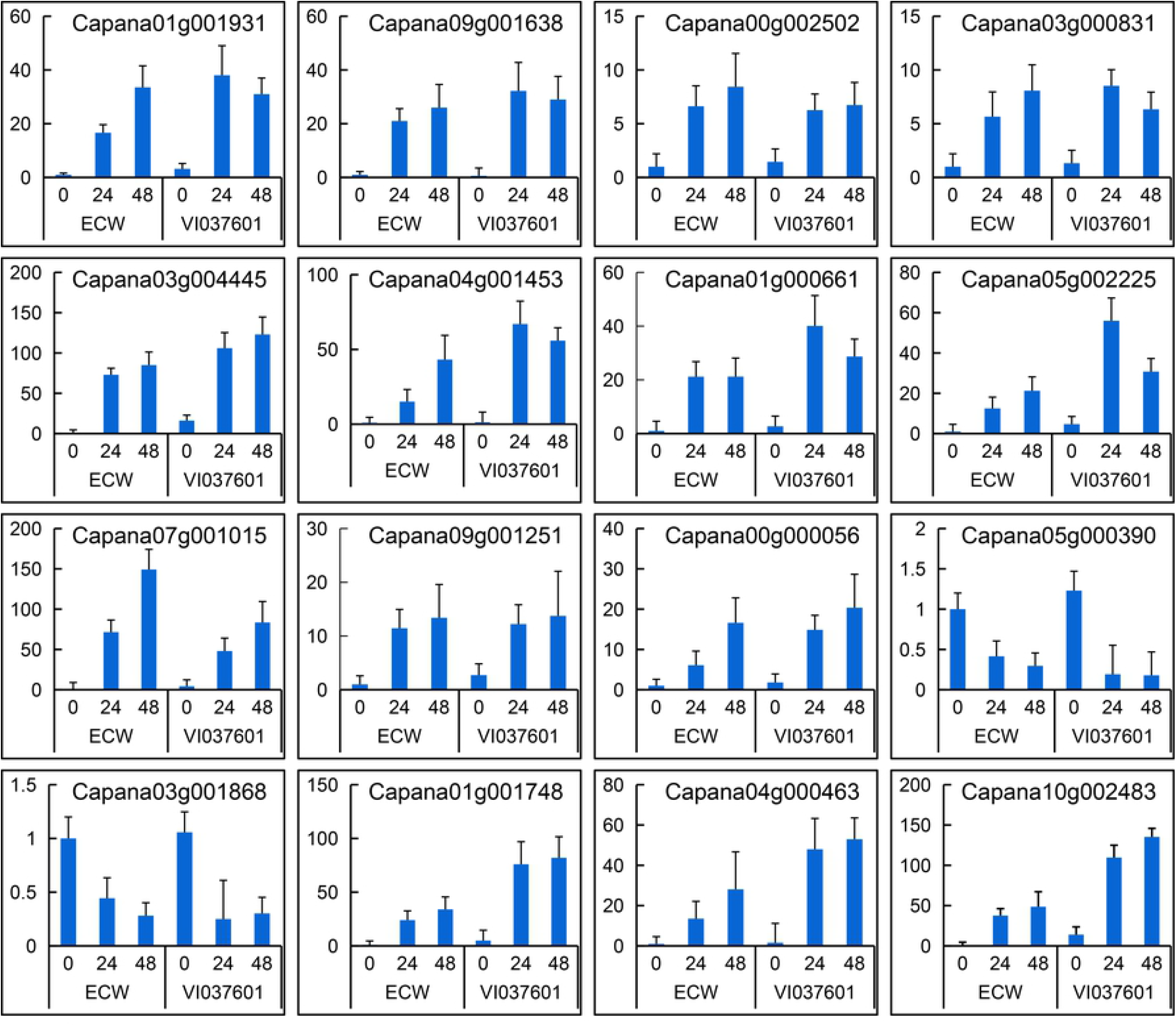
qRT-PCR based validation of DEGs in response to *Xcv* inoculation at different time intervals. 0 hpi represented the time points at 0 hpi in ECW and VI037601 with the mock inoculation of sterile water. 24 hpi and 48 hpi represented the time points at 24 hours and 48 hours post *Xcv* inoculation in ECW and VI037601, respectively. The data were normalized by using ubiquitin *CaUbi3* as an internal reference. Data were represented as mean ± SD for three biological replicates.

## Discussion

Plants are exposed to a myriad of pathogenic microorganisms during their lifespan, including bacteria, fungi, viruses and nematodes, which all try to acquire nutrients from the host plant for their advantage [38]. BS caused by *Xcv* is a very serious global disease, which has caused enormous yield and economic losses in pepper production, especially in regions with a warm and humid climate. In response to bacterial attack, plants activate a wide array of defenses which can reduce bacteria damage, such as mitogen-activated protein kinase (MPK) cascades, the induction of defense-related signaling molecule biosynthesis [39]. The *Capsicum annuum* L. cultivar VI037601 was considered as a resistant source to BS due to hypersensitive response reaction to *Xcv* strain 23-1, while cultivar ECW was susceptible to *Xcv* infection, which was also confirmed in this study (Fig 1). Yet, the molecular mechanisms to respond *Xcv* infection between VI037601 and ECW were not clear. In this study, RNA-seq technique was adapted to identify the DEGs associated with disease response during the race *Xcv* strain 23-1 infection in the leaves of ECW and VI037601. An average of 6.9 G clean reads was obtained from RNA-seq in each library, and at least 71.94% of which were matched to the reference genome sequence Zunla-1 (Table 1; Table S1). Besides that, we detected 52,043 transcript genes with 16,705 novel genes in the eighteen libraries (Table S2). These datum indicated that the sequencing depth was sufficient for the transcritpme coverage. Besides, the good correlation between qRT-PCR and RNA-seq data also demonstrated the reliability of the RNA-seq analysis (Fig 6). Therefore, the analysis of RNA-seq data would provide new insights into the response of pepper to *Xcv* inoculation.

Plants have an innate immunity system to defend themselves against pathogens by a number of mechanisms, such as hypersensitive response (HR), induction of genes encoding PR, induced biosynthesis of secondary metabolites [40-41]. DEGs in ECW and VI037601 post *Xcv* inoculation and KEGG enrichment analysis demonstrated that many biological processes were influenced by pathogen infection, including amino sugar and nucleotide sugar metabolism (ko00520), sesquiterpenoid and triterpenoid biosynthesis (ko00909), MAPK signaling pathway (ko04016), plant-pathogen interaction (ko04626) and plant hormone signal transduction (ko04075) (Table 2; Table S7). Furthermore, many DEGs were enriched in oxidoreductase activity (GO:0016684), acting on peroxide as acceptor (GO:0016684), peroxidase activity (GO:0004601), chitinase activity (GO:0004568), chitin binding (GO:0008061), amino sugar catabolic process (GO:0046348), amino sugar metabolic process (GO:0006040), aminoglycan catabolic process (GO:0006026), chitin catabolic process (GO:0006032), and chitin metabolic process (GO:0006030) (Fig 3; Table S6). Similar results were obtained in the previous study of tomato response to infection by *Xanthomonas* perforans Race 3 [24]. The expression of these defense response genes induced the synthesis of secondary metabolites, which could inhibit the spread of *Xcv* in peppers.

In plants, the hypersensitive response (HR) is a form of programmed cell death (PCD) at the site of pathogen infection, which is closely related to active resistance [42]. Previous studies showed that *Bs2, Bs3* and *Bs4* were only expressed in BS resistant pepper post *Xcv* inoculation, which could trigger HR [15,16]. Here, transcriptome profiling analysis results showed that *bs2* (Capana09g000438) and *bs3* (Capana02g001306) were not/hardly expressed in pepper leaves before and after *Xcv* inoculation. Moerover, the expression of *bs4* (XLOC_032013), as well as the homologs of *bs2* and *bs3*, did not change significantly. However, many proteins kinases/enzymes encoded by DEGs were involved in defense-related gene induction and innate immunity, such as those that activate genes coding for the receptor-like kinases (RLKs), NAC TFs, WRKY TFs, pathogenesis-related protein and chitinase, as reported previously [21,43]. In this study, among 273 DEGs were specific differentially expressed in VI037601 post *Xcv* inoculation, many protein kinase, oxidordeuctase, TFs and uncharacterized proteins were identified, such as Capana10g000475 (receptor-like protein kinase), Capana09g000463 (transcription factor), Capana02g003648 (peroxidase) and XLOC_024869 (uncharacterized protein) (Table S5; Table S8). Furthermore, Capana09g000319 (aldehyde dehydrogenase) and Capana09g000326 (glycosyltransferase) were specifically expressed in VI037601, and the expression of which was significantly up-regulated after *Xcv* inoculation (Table S5), indicating that they might play an important role in response to *Xcv* infection. Receptor-like kinases act as pattern-recognition receptors, which recognize pathogens as the first layer of inducible defense [44]. Our findings also showed that many receptor-like kinases were significantly differentially expressed in ECW and VI037601, such as G-type lectin S-receptor-like serine/threonine-protein kinase (Capana07g002260) and LRR receptor-like serine/threonine-protein kinase (Capana03g000831), which were up regulated in ECW and VI037601 post *Xcv* inoculation (Fig 4; Table S5; Table S8). RLKs were important signaling components that played key roles in adapting to numerous biotic and abiotic stresses as well as in regulating plant growth and development [39,45]. Generally, the mitogen-activated protein kinase (MPK) cascades were initiated by the stimulated receptors. After a series of cascades reactions, activated MPKs phosphorylated their substrates, most of which were enzymes and transcription factors, thereby triggering downstream responses [46]. WRKY TFs as the substrates of MPKs can be regulated by MPKs at transcriptional and/or post-translational levels [47-49]. For instance, OsWRKY53 was activated by OsMPK3 and OsMPK6 through transcriptional induction and phosphorylation in the process of pathogen infection, thereby enhancing rice resistance to pathogens [46,50]. Here, eight WRKY TFs were also up-regulated in ECW and VI037601 post *Xcv* inoculation (Fig 5; Table S5; Table S8; Table S9), which might be induced by MPKs. Up-regulated expression of these WRKY TFs could activate downstream disease response genes or hormones pathway-related genes to protect against pathogen infection [37]. Besides, many studies showed that TFs that contain the NAC domain played pivotal roles in the regulation of the transcriptional reprogramming associated with plant stress responses, such as abiotic stress response and pathogen defense [51]. These NAC proteins might positively regulate plant defense responses by activating PR genes, including HR, and cell death at the infection site, such as ATAF1, which positively regulated penetration resistance to biotrophic fungus *Blumeria graminis* f.sp. *hordei* (*Bfh*) [52]. *OsNAC6, ONAC066, ONAC122, ONAC131, OsNAC111*, and *OsNAC4* have been validated to be involved in defense responses against pathogen attack [53-56]. MYB TFs also played important roles in response to pathogen infection. Overexpression of *SmMYB44* in eggplant increased the resistance to bacterial wilt [57]. Here, two MYB TFs were differentially expressed in VI037601, while three MYB TFs and six NAC TFs were up-regulated in VI037601 and ECW post *Xcv* inoculation, which might play pivotal roles in pathogen defense.

Moreover, many genes in the “plant hormone signal transduction” pathway were up or down regulated during symptom development. Plant resistance to biotrophic pathogens is positively regulated by ethylene and is negatively regulated by the auxin signal transduction pathway [58,59]. Ethylene-responsive transcription factors mediated disease resistance was demonstrated in Arabidopsis and tomato against *Botrytis cinerea* and *Ralstonia solanacearum*, respectively [60-62]. Here, six ERFs were specific differentially exoressed in VI037601 post *Xcv* inoculation and six ERFs were up regulated in ECW and VI037601 post *Xcv* inoculation (Fig 5; Table S8). Besides that, the overlapping DEGs associated with auxin-responsive genes were down regulated in ECW and VI037601 post *Xcv* inoculation, such as Capana03g004567 and Capana03g000299 (Fig 5; Table S8). Thus, these DEGs might implicate their roles in the regulation of transcriptional reprogramming associated with the early response to *Xcv* in pepper.

## Conclusions

This is the first study that examined global transcriptional changes in pepper plants infected with *Xcv*. In this study, 4,794 DEGs were identified in ECW and VI037601 post *Xcv* inoculation, which were mainly enriched in single-organism metabolic process (GO:0044710), oxidation-reduction process (GO:0055114), response to stress (GO:0006950), defense response (GO:0006952), and secondary metabolic process (GO:0019748). DEGs that belonged to defense response, were explored, such as protein kinase genes, oxidoreductase, and TFs. Most of which responded faster in VI037601 than that in ECW post *Xcv* inoculation, indicating that these DEGs might play vital roles in response to BS disease at early infection stage. This study provides important information to elucidate the defense molecular mechanisms of pepper during *Xcv* infection.

## Supplementary information

**Table S1: Statistic analysis of pepper clean reads in 18 libraries for RNA-seq.**

**Table S2: Detailed list of novel genes**.

**Table S3: Genes FPKM value, annotation, and function enrichment**.

**Table S4: Detailed list of DEGs in ECW and VI037601 at 24 hpi and 48 hpi relative to 0 hpi**.

**Table S5: Detailed list of specific differentially expressed genes in VI037601 post *Xcv* inoculation and overlapping DEGs in ECW and VI037601 at 24 hpi and 48 hpi relative to 0 hpi**.

**Table S6: Summary of GO terms related to the differentially expressed genes in ECW and VI037601 post *Xcv* inoculation**.

**Table S7: Significantly enriched KEGG pathway in ECW and VI037601 after infection of *Xcv*.**

**Table S8: DEGs related to BS disease resistance in pepper**.

**Table S9: DEGs involved in transcription factor families in pepper**.

**Table S10: qRT-PCR primers for the validation of RNA-Seq data**.

## Author contributions

**Conceptualization:** Shenghua Gao, Fei Wang, Chunhai Jiao, Minghua Yao.

**Data curation:** Shenghua Gao, Fei Wang.

**Formal analysis:** Shenghua Gao, Juntawong Niran, Ning Li, Yanxu Yin, Chuying Yu.

**Funding acquisition:** Shenghua Gao, Chunhai Jiao, Minghua Yao.

**Investigation:** Fei Wang, Juntawong Niran, Ning Li, Yanxu Yin, Chuying Yu.

**Supervision:** Chunhai Jiao, Minghua Yao.

**Writing - original draft:** Shenghua Gao, Fei Wang, Chunhai Jiao, Minghua Yao.

**Writing - review & editing:** Shenghua Gao, Fei Wang, Chunhai Jiao, Minghua Yao.

## Acknowledgments

We thank Professors Gongyou Chen of Shanghai Jiao Tong University and Meng Yuan of Huazhong Agricultural University for kindly providing *Xanthomonas campestris* pv. *vesicatoria* (*Xcv*). Thanks also to Frasergen Bioinformatics Co., Ltd for RNA-Sequencing and Professor Changxian Yang of Huazhong Agricultural University for correcting the English in this paper.

## References

1. Truong HTH, Kim KT, Kim S, Cho MC, Kim HR, Woo JG. Development of gene-based markers for the *Bs2* bacterial spot resistance gene for marker-assisted selection in pepper (*Capsicum* spp.). Hort Environ Biotechnol. 2011; 52(1): 65–73. doi: 10.1007/s13580-011-0142-4

2. Baek W, Lim S, Lee SC. Identification and functional characterization of the pepper *CaDRT1* gene involved in the ABA-mediated drought stress response. Plant Mol Biol. 2016; 91(1-2): 149–160. doi: 10.1007/s11103-016-0451-1

3. Bulle M, Yarra R, Abbagani S. Enhanced salinity stress tolerance in transgenic chilli pepper (*Capsicum annuum* L.) plants overexpressing the wheat antiporter (*TaNHX2*) gene. Mol Breeding. 2016; 36(4): 36. doi: 10.1007/s11032-016-0451-5

4. Mishra R, Mohanty JN, Chand SK, Joshi RK. Can-miRn37a mediated suppression of ethylene response factors enhances the resistance of chilli against anthracnose pathogen *Colletotrichum truncatum* L. Plant Sci. 2018; 267: 135–147. doi: 10.1016/j.plantsci.2017.12.001

5. Tai T, Dahlbeck D, Stall D, Peleman J, Staskawicz BJ. High-resolution genetic and physical mapping of the region containing the *Bs2* resistance gene of pepper. Theor Appl Genet. 1999; 9: 1201–1206. doi: 10.1007/s001220051325

6. Thieme F, Koebnik R, Bekel T, Berger C, Boch J, Büttner D, et al. Insights into genome plasticity and pathogenicity of the plant pathogenic bacterium *Xanthomonas campestris* pv. *vesicatoria* revealed by the complete genome sequence. J Bacteriol. 2005; 187(21): 7254–7266. doi: 10.1128/JB.187.21.7254-7266.2005

7. Stall RE, Jones JB, Minsavage GV. Durability of Resistance in Tomato and Pepper to *Xanthomonads* Causing Bacterial Spot. Annu Rev Phytopathol. 2009; 47(1): 265–284. doi: 10.1146/annurev-phyto-080508-081752

8. Jones JB, Stall RE, Bouzar H. Diversity among *Xanthomonads* pathogenic on pepper and tomato. Annu Rev Phytopathol. 1998; 36: 41–58. doi: 10.1146/annurev.phyto.36.1.41

9. Ibrahim Y, Al-saleh M. First report of bacterial spot caused by *Xanthomonas campestris* pv. *vesicatoria* on sweet pepper (*Capsicum annuum* L.) in Saudi Arabia. Plant Dis. 2012; 96(11):1690–1690. doi: 10.1094/PDIS-04-12-0354-PDN

10. Jibrin MO, Timilsina S, Potnis N, Minsavage GV, Shenge KC, Akpa AD, et al. First report of *Xanthomonas euvesicatoria* causing bacterial spot disease in pepper in Northwestern Nigeria. Plant Dis. 2014; 98(10): 1426–1427. doi: 10.1094/PDIS-06-14-0586-PDN

11. Griffin K, Gambley C, Brown P, Li Y. Copper-tolerance in *Pseudomonas syringae* pv. tomato and *Xanthomonas* spp and the control of diseases associated with these pathogens in tomato and pepper. A systematic literature review. Crop Prot. 2017; 96: 144–150. doi: 10.1016/j.cropro.2017.02.008

12. Jones JB, Minsavage GV, Roberts PD, Johnson RR, Kousik CS, Subramanian S, Stall RE. A non-hypersensitive resistance in pepper to the bacterial spot pathogen is associated with two recessive genes. Phytopathology. 2002; 92(3): 273–277. doi: 10.1094/PHYTO.2002.92.3.273

13. Pierre M, Noël L, Lahaye T, Ballvora J, Veuskens J, Ganal M, Bonas U. High-resolution genetic mapping of the pepper resistance locus *Bs3* governing recognition of the *Xanthomonas campestris* pv *vesicatoria* AvrBs3 protein. Theor Appl Genet. 2000; 101(1-2): 255–263. doi: 10.1007/s001220051477

14. Tai T, Staskawicz BJ. Construction of a yeast artificial chromosome library of pepper (*Capsicum annuum* L.) and identification of clones from the *Bs2* resistance locus. Theor Appl Genet. 2000; 100(1): 112–117. doi: 10.1007/s001220050016

15. Römer P, Hahn S, Jordan T, Strauß T, Bonas U, Lahaye T. Plant pathogen recognition mediated by promoter activation of the pepper *Bs3* resistance gene. Science. 2007; 318: 645–648. doi: 10.1126/science.1144958

16. Wang J, Zeng X, Tian D, Yang X, Wang L, Yin Z. The pepper Bs4C proteins are localized to the endoplasmic reticulum (ER) membrane and confer disease resistance to bacterial blight in transgenic rice. Mol Plant Pathol. 2018; 19(8): 2025–2035. doi: 10.1111/mpp.12684

17. Garg T, Mallikarjuna BP, Thudi M, Samineni S, Singh S, Sandhu JS, et al. Identification of QTLs for resistance to Fusarium wilt and Ascochyta blight in a recombinant inbred population of chickpea (*Cicer arietinum* L.). Euphytica. 2018; 214: 45. doi: 10.1007/s10681-018-2125-3

18. Chen H, Lu C, Jiang H, Peng J. Global transcriptome analysis reveals distinct aluminum-tolerance pathways in the Al-accumulating species Hydrangea macrophylla and marker identification. PLoS One. 2015; 10(12): e0144927. doi: 10.1371/journal.pone.0144927

19. Li T, Xu X, Li Y, Wang H, Li Z, Li Z. Comparative transcriptome analysis reveals differential transcription in heat-susceptible and heat-tolerant pepper (*Capsicum annum* L.) cultivars under heat stress. J Plant Biol. 2015; 58(6): 411–424. doi: 10.1007/s12374-015-0423-z

20. Ou L, Liu Z, Zhang Z, Wei G, Zhang Y, Kang L, et al. Noncoding and coding transcriptome analysis reveals the regulation roles of long noncoding RNAs in fruit development of hot pepper (*Capsicum annuum* L.). Plant Growth Regul. 2017; 83(1): 141–156. doi: 10.1007/s10725-017-0290-3

21. Zhu C, Li X, Zheng J. Transcriptome profiling using Illumina-and SMRT-based RNA-seq of hot pepper for in-depth understanding of genes involved in CMV infection. Gene. 2018; 666: 123–133. doi: 10.1016/j.gene.2018.05.004

22. Zuo J, Wang Y, Zhu B, Luo Y, Wang Q, Gao L. Analysis of the coding and non-coding RNA transcriptomes in response to bell pepper chilling. Int J Mol Sci. 2018; 19(7): 2001. doi: 10.3390/ijms19072001

23. Balaji V, Gibly A, Debbie P, Sessa G. Transcriptional analysis of the tomato resistance response triggered by recognition of the *Xanthomonas* type III effector AvrXv3. Funct Integr Genomic. 2007; 7(4): 305–316. doi: 10.1007/s10142-007-0050-y

24. Du H, Wang Y, Yang J, Yang W. Comparative Transcriptome analysis of resistant and susceptible tomato lines in response to infection by *Xanthomonas* perforans Race T3. Front Plant Sci. 2015; doi: 10.3389/fpls.2015.01173

25. Huang R, Hui S, Zhang M, Li P, Xiao J, Li X, et al. A conserved basal transcription factor is required for the function of diverse TAL effectors in multiple plant hosts. Front Plant Sci. 2017; doi: 10.3389/fpls.2017.01919.

26. Trapnell C, Roberts A, Goff L, Pertea G, Kim D, Kelley DR. et al. Differential gene and transcript expression analysis of RNA-Seq experiments with TopHat and Cufflinks. Nat Protoc. 2012; 7(3): 562–578. doi: 10.1038/nprot.2012.016

27. Trapnell C, Williams BA, Pertea G, Mortazavi A, Kwan G, van Baren MJ, et al. Transcript assembly and quantification by RNA-Seq reveals unannotated transcripts and isoform switching during cell differentiation. Nat Biotechnol. 2010; 28(5): 511–515. doi: 10.1038/nbt.1621

28. Anders S, Huber W. Differential expression analysis for sequence count data. Genome Biol. 2010; 11(10): R106. doi: 10.1186/gb-2010-11-10-r106

29. Conesa A, Götz S. Blast2GO: A comprehensive suite for functional analysis in plant genomics. Int J Plant Genomics. 2008; 1–12. doi: 10.1155/2008/619832

30. Jin J, Tian F, Yang D.C, Meng Y.Q, Kong L, Luo J, Gao G. PlantTFDB 4.0: toward a central hub for transcription factors and regulatory interactions in plants. Nucleic Acids Res. 2017; 45 (D1): D1040–5. doi: 10.1093/nar/gkw1328

31. Wan H, Yuan W, Ruan M, Ye Q, Wang R, Li Z, et al. Identification of reference genes for reverse transcription quantitative real-time PCR normalization in pepper (*Capsicum annuum* L.). Biochem Bioph Res Co. 2011; 416(1-2): 24–30. doi: 10.1016/j.bbrc.2011.10.105

32. Schmittgen TD, Livak KJ. Analyzing real-time PCR data by the comparative CT method. Nat Protoc. 2008; 3(6): 1101–1108. doi: 10.1038/nprot.2008.73>

33. Trapnell C, Pachter L, Salzberg SL. TopHat: discovering splice junctions with RNA-Seq. Bioinformatics. 2009; 25(9): 1105–1111. doi: 10.1093/bioinformatics/btp120

34. Ashburner M, Ball CA, Blake JA, Botstein D, Butler H, Cherry JM, et al. Gene ontology: tool for the unification of biology. The Gene Ontology Consortium. Nat Genet. 2000; 25(1): 25–29. doi: 10.1038/75556

35. Kanehisa M, Goto S, Kawashima S, Okuno Y, Hattori M. The KEGG resource for deciphering the genome. Nucleic Acids Res. 2004; 32: D277–280. doi: 10.1093/nar/gkh063

36. Xie C, Mao X, Huang J, Ding Y, Wu J, Dong S, Kong L, Gao G, Li CY, Wei L. KOBAS 2.0: a web server for annotation and identification of enriched pathways and diseases. Nucleic Acids Res. 2011; 39: W316–322. doi: 10.1093/nar/gkr483

37. Tariq R, Wang C, Qin T, Xu F, Tang Y, Gao Y, Ji Z, Zhao K. Comparative Transcriptome profiling of rice near-isogenic line carrying *Xa23* under infection of *Xanthomonas oryzae* pv. *oryzae*. Int J Mol Sci. 2018; 19(3): 717. doi: 10.3390/ijms19030717

38. Lee DS, Kim YC, Kwon SJ, Ryu CM, Park OK. The Arabidopsis cysteine-rich receptor-like kinase CRK36 regulates immunity through interaction with the cytoplasmic kinase BIK1. Front Plant Sci. 2017; doi: 10.3389/fpls.2017.01856.

39. Boller T, Felix G. A renaissance of elicitors: perception of microbe-associated molecular patterns and danger signals by pattern-recognition receptors. Annu Rev Plant Biol. 2009; 60(1): 379–406. doi: 10.1146/annurev.arplant.57.032905.105346

40. Muthamilarasan M, Prasad M. Plant innate immunity: An updated insight into defense mechanism. J Biosciences. 2013; 38(2): 433–449. doi: 10.1007/s12038-013-9302-2

41. Balakireva A, Zamyatnin A. Indispensable Role of Proteases in Plant Innate Immunity. Int J Mol Sci. 2018; 19(2): 629. doi: 10.3390/ijms19020629

42. Olukolu BA, Negeri A, Dhawan R, Venkata BP, Sharma P, Garg A, Gachomo E, Marla S, Chu K, Hasan A, Ji J, Chintamanani S, Green J, Shyu C, Wisser R, Holland J, Johal G, Balint-Kurti P. A connected set of genes associated with programmed cell death implicated in controlling the Hypersensitive response in maize. Genetics. 2012; 193(2): 609–620. doi: 10.1534/genetics.112.147595

43. Matsubayashi Y, Ogawa M, Morita A, Sakagami Y. An LRR receptor kinase involved in perception of a peptide plant hormone, phytosulfokine. Science. 2002; 296(5572): 1470–1472. doi: 10.1126/science.1069607

44. Tripathi L, Tripathi JN, Shah T, Muiruri KS, Katari M. Molecular basis of disease resistance in banana progenitor musa balbisiana against *Xanthomonas campestris* pv. *musacearum*. Sci Rep. 2019; 9(1). doi: 10.1038/s41598-019-43421-1

45. Ou Y, Lu X, Zi Q, Xun Q, Zhang J, Wu Y, et al. RGF1 INSENSITIVE 1 to 5, a group of LRR receptor-like kinases, are essential for the perception of root meristem growth factor 1 in *Arabidopsis thaliana*. Cell Res. 2016; 26(6): 686–698. doi: 10.1038/cr.2016.63

46. Hu L, Ye M, Kuai P, Ye M, Erb M, Lou Y. OsLRR-RLK1, an early responsive leucine-rich repeat receptor-like kinase, initiates rice defense responses against a chewing herbivore. New Phytol. 2018; 219(3): 1097–1111. doi: 10.1111/nph.15247

47. Ishihama N, Yoshioka H. Post-translational regulation of WRKY transcription factors in plant immunity. Curr Opin Plant Biol. 2012; 15: 431–437. doi: 10.1016/j.pbi.2012.02.003

48. Chi Y, Yan Y, Zhou Y, Zhou J, Fan B, Yu JQ, Chen Z. Protein-protein interactions in the regulation of WRKY transcription factors. Mol Plant. 2013; 6: 287–300. doi: 10.1093/mp/sst026

49. Li R, Zhang J, Li J, Zhou G, Wang Q, Bian W, et al. Prioritizing plant defence over growth through WRKY regulation facilitates infestation by non-target herbivores. ELife. 2015; doi: 10.7554/eLife.04805.

50. Chujo T, Miyamoto K, Ogawa S, Masuda Y, Shimizu T, Kishi-Kaboshi M, et al. Overexpression of phosphomimic mutated *OsWRKY53* leads to enhanced blast resistance in rice. PloS One. 2014; 9: e98737. doi: 10.1371/journal.pone.0098737

51. Nuruzzaman M, Sharoni AM, Kikuchi S. Roles of NAC transcription factors in the regulation of biotic and abiotic stress responses in plants. Front Microbiol. 2013; 4: 248. doi: 10.3389/fmicb.2013.00248

52. Jensen MK, Rung JH, Gregersen PL, Gjetting T, Fuglsang AT, Hansen M, et al. The *HvNAC6* transcription factor: a positive regulator of penetration resistance in Barley and Arabidopsis. Plant Mol Biol. 2007; 65(1-2): 137–150. doi: 10.1007/s11103-007-9204-5

53. Nakashima K, Tran LS, Van Nguyen D, Fujita M, Maruyama K, Todaka D, Ito Y, Hayashi N, Shinozaki K, Yamaguchi-Shinozaki K. Functional analysis of a NAC-type transcription factor *OsNAC6* involved in abiotic and biotic stress-responsive gene expression in rice. Plant J. 2007; 51: 617–630. doi: 10.1111/j.1365-313x.2007.03168.x

54. Kaneda T, Taga Y, Takai R, Iwano M, Matsui H, Takayama S, Isogai A, Che FS. The transcription factor *OsNAC4* is a key positive regulator of plant hypersensitive cell death. EMBO J. 2009; 28: 926–936. doi: 10.1038/emboj.2009.39

55. Sun L, Zhang H, Li D, Huang L, Hong Y, Ding XS. Nelson RS, Zhou X, Song F. Functions of rice NAC transcriptional factors, *ONAC122* and *ONAC131*, in defense responses against Magnaporthe grisea. Plant Mol Biol. 2013; 81: 41–56. doi: 10.1007/s11103-012-9981-3

56. Liu Q, Yan S, Huang W, Yang J, Dong J, Zhang S, Zhao J, Yang T, Mao X, Zhu X, Liu B. NAC transcription factor *ONAC066* positively regulates disease resistance by suppressing the ABA signaling pathway in rice. Plant Mol Biol. 2018; doi: 10.1007/s11103-018-0768-z

57. Qiu Z, Yan S, Xia B, Jiang J, Yu B, Lei J, Chen C, Lin Chen L, Yang Y, Wang Y, Tian S, Cao B. The eggplant transcription factor *MYB44* enhances resistance to bacterial wilt by activating the expression of spermidine synthase. J Exp Bot. 2019; doi: 10.1093/jxb/erz259

58. Ferrari S, Galletti R, Pontiggia D, Manfredini C, Lionetti V, Bellincampi D, et al. Transgenic expression of a fungal endo-polygalacturonase increases plant resistance to pathogens and reduces auxin sensitivity. Plant Physiol. 2008; 146(2): 669–681. doi: 10.1104/pp.107.109686

59. Sun X, Yu G, Li J, Liu J, Wang X, Zhu G, Zhang X, Pan H. *AcERF2*, an ethylene-responsive factor of Atriplex canescens, positively modulates osmotic and disease resistance in *Arabidopsis thaliana*. Plant Sci. 2018; 274: 32–43. doi: 10.1016/j.plantsci.2018.05.004

60. Su RC, Cheng CP, Sanjaya, You SJ, Hsieh TH, Chao TC, Chan MT. Tomato RAV transcription factor is a pivotal modulator involved in the AP2/EREBP-mediated defense pathway. Plant Physiol. 2011; 156, 213–27. doi: 10.1104/pp.111.174268

61. Zhao Y, Wei T, Yin KQ, Chen Z, Gu H, Qu L, Qin G. Arabidopsis RAP2.2 plays an important role in plant resistance to *Botrytis cinerea* and ethylene responses. New Phytol. 2012; 195: 450–60. doi: 10.1111/j.1469-8137.2012.04160.x

62. Moffat CS, Ingle RA, Wathugala DL, Saunders NJ, Knight H, Knight MR. ERF5 and ERF6 play redundant roles as positive regulators of JA/Et-mediated defense against *Botrytis cinerea* in *Arabidopsis*. PLoS One. 2012; 7: e35995, doi: 10.1371/journal.pone.0035995

